# The glycosyltransferase ALG3 is an AKT substrate that regulates protein N-glycosylation

**DOI:** 10.1101/2025.04.01.646556

**Authors:** Adrija J Navarro-Traxler, Laura Ghisolfi, Evan C Lien, Alex Toker

## Abstract

The PI3K/AKT signaling pathway is frequently dysregulated in cancer and controls key cellular processes such as survival, proliferation, metabolism and growth. Protein glycosylation is essential for proper protein folding and is also often deregulated in cancer. Cancer cells depend on increased protein folding to sustain oncogene-driven proliferation rates. The N-glycosyltransferase asparagine-linked glycosylation 3 homolog (ALG3), a rate-limiting enzyme during glycan biosynthesis, catalyzes the addition of the first mannose to glycans in an alpha-1,3 linkage. Here we show that ALG3 is phosphorylated downstream of the PI3K/AKT pathway in both growth factor-stimulated cells and PI3K/AKT hyperactive cancer cells. AKT directly phosphorylates ALG3 in the amino terminal region at Ser11/Ser13. CRISPR/Cas9-mediated depletion of ALG3 leads to improper glycan formation and induction of endoplasmic reticulum stress, the unfolded protein response, and impaired cell proliferation. Phosphorylation of ALG3 at Ser11/Ser13 is required for glycosylation of cell surface receptors EGFR, HER3 and E-cadherin. These findings provide a direct link between PI3K/AKT signaling and protein glycosylation in cancer cells.

## INTRODUCTION

The phosphoinositide 3-kinase (PI3K)/AKT pathway is a critical signaling network that controls cell growth, proliferation, metabolism and survival. In response to extracellular stimuli including growth factors and hormones, activation of PI3K leads to synthesis of PIP3 and PI(3,4)P2, lipid second messengers that recruit downstream effectors to promote signal relay^1–3^. Of these, the Ser/Thr protein kinase AKT is a major transducer of the PI3K signal that regulates a diverse array of substrates including protein and lipid kinases, transcription factors, small G proteins, metabolic enzymes, E3 ubiquitin ligases and cell cycle regulators^1^. AKT is a basophilic-directed protein kinase that phosphorylates Ser/Thr residues within a defined consensus motif of RXRXXs/t-Φ (where X is any amino acid, and Φ is any bulky hydrophobic residue)^4^. Over 200 AKT substrates that harbor this consensus motif have been identified, whereby phosphorylation regulates enzymatic activity, cellular localization and protein-protein interactions, leading to functional consequences in normal physiology and pathophysiology^1,4,5^.

PI3K/AKT is one of the most frequently genetically-dysregulated pathways in human cancers, due to somatic mutations and amplification of oncogenes such as *PIK3CA* (encoding the p110α catalytic subunit of class I PI3K) and *AKT1*, as well as inactivation of tumor suppressors such as phosphatase and tensin homolog (*PTEN*) and inositol polyphosphate 4-phosphatase type II (*INPP4B*)^6,7^. Mechanistically, PI3K/AKT signaling modulates cellular anabolic metabolism through the phosphorylation and regulation of rate-limiting metabolic enzymes involved in nucleotide, protein and lipid biosynthesis^8^. PI3K/AKT also controls glucose and glycogen metabolism, influencing glycolysis at the transcriptional and post-translational level^9–12^. By contrast, a major role for PI3K/AKT signaling in the control of carbohydrate metabolism has not been described^13^. Glycosylation, the process by which complex carbohydrates, or glycans, are covalently added to proteins, sugars, and lipids, is required for all aspects of normal cellular physiology and homeostasis^14–16^. N-glycosylation, in which glycans are added to asparagine residues, is functionally critical for cell-surface receptors, secreted proteins, lysosomal enzymes, immunoglobulins, tumor antigens and integrins^14,15,17–22^. N-glycosylation is altered during the metastatic transition in cancer, the epithelial to mesenchymal transition, and during anti-tumor immune responses^23,24^. Moreover, genes whose protein products participate in glycan synthesis, degradation and modification are dysregulated in cancer, leading to aberrant glycan synthesis, or incomplete or truncated glycan structures^19,25–35^.

N-glycosylation is initiated in the endoplasmic reticulum (ER) and is necessary for proper protein folding and ER homeostasis^36–38^. In the ER, glycosyltransferases catalyze the addition of specific carbohydrate moieties to a nascent glycan chain, with dolichol serving as a lipid carrier^13,37–40^. Of these, the asparagine-linked glycosylation 3 homolog alpha-1,3-mannosyltransferase (ALG3), catalyzes the addition of the first Dol-P-Man derived mannose to man5GlcNAc2-PP-Dol in an alpha-1,3 linkage^41,42^.

ALG3, an evolutionarily-conserved glycosyltransferase, has been functionally characterized in plants, fungi, yeast, and fruit flies. In humans, compound heterozygous mutations in *ALG3* lead to congenital disorders of N-linked glycosylation (ALG3-CDG)^43–47^. Patients with this rare disorder display severe neurological dysfunction, developmental delays, intellectual disabilities and failure to thrive^43–47^. Homozygous mutations in *ALG3* are rare, suggesting these may be lethal in the context of embryonic development^43^. Depletion of ALG3 in cells leads to improper glycan formation and induction of ER stress^46,48,49^. In human cancers, *ALG3* is amplified in a variety of lineages, including lung, breast, ovarian, and esophageal cancers^50–52^. Studies have also indicated that ALG3 expression is correlated with poor outcome in breast cancer^51,52^. Whether ALG3 or other N-linked glycosyltransferases are regulated by classical signaling pathways by post-translational modifications (PTMs) has not been determined.

Here, we report that ALG3 is a target of PI3K signaling and is directly phosphorylated by AKT. Depletion of ALG3 induces ER stress and the unfolded protein response (UPR) with a concomitant deregulation of glycoproteins and reduced cell proliferation in breast cancer cells.

## RESULTS

### ALG3 is an AKT substrate downstream of PI3K

To determine if glycosyltransferases represent a distinct class of AKT downstream targets, we screened the PhosphoSitePlus database (www.phosphosite.org) for proteins that have been mapped by global mass spectrometry phosphoproteomic studies and which contain the minimal AKT consensus motif RXRXXs/t^53^. This analysis revealed a number of glycosyltransferases which harbor PTMs including Ser/Thr/Tyr phosphorylation, Lys/Arg mono-methylation, Lys ubiquitylation, Lys acetylation, Lys succinylation, Asn N-glycosylation, Ser/Thr O-GlcNAc and Arg di-methylation. Within this group, ALG3 contains two high-quality basophilic phosphorylation motifs at Ser11/Ser13 that conform to the optimal AKT consensus motif (**Fig. 1A**)^5^.

**Figure 1.**
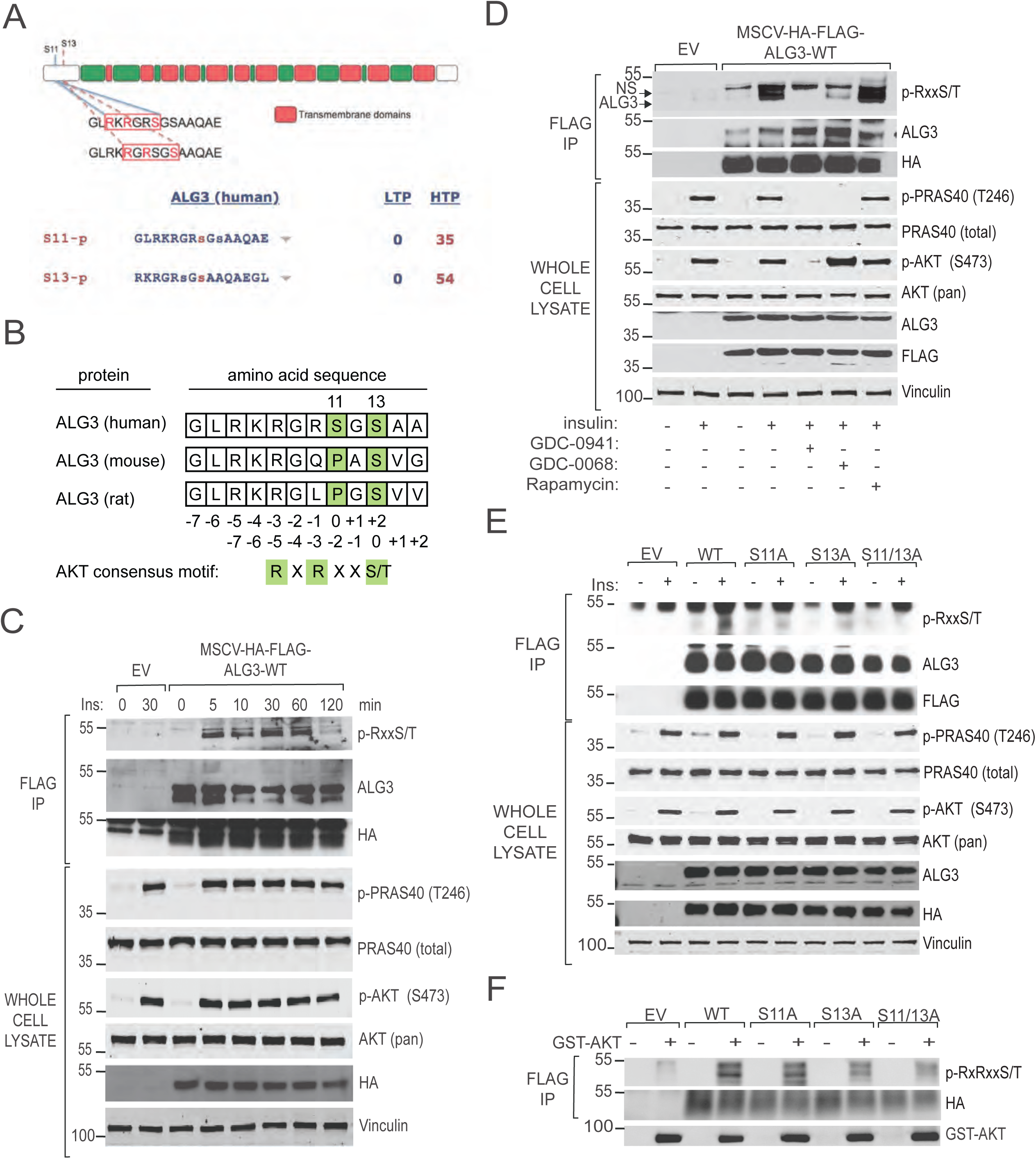
ALG3 is an AKT substrate. (A) ALG3 contains the AKT substrate motif RXRXX-S/T at Ser11 and Ser13. (B) ALG3 phosphosite conservation in mammals. Serum-starved MCF10A cells expressing EV or HA-FLAG-ALG3 (C) were treated with 100 nM insulin for the indicated times (D) in the presence and absence of 15 min inhibitor pretreatment: 2 μM GDC-0941, 2 μM GDC-0068, or 20 nM rapamycin. FLAG immunoprecipitants from cell lysates were immunoblotted. NS arrow denotes a non-significant background band. The arrow denotes ALG3 protein. (E) Serum-starved MCF10A cells expressing EV, HA-FLAG-ALG3-WT, -S11A, -S13A or -S11/13A were treated with 100 nM insulin. FLAG immunoprecipitants were immunoblotted. (F) EV, HA-FLAG-ALG3-WT, -S11A, -S13A and -S11/13A (containing an additional 11 amino acids appended to the C-terminus of ALG3 with no phosphorylation sites) were immunoprecipitated from serum starved MCF10A cells and incubated with active GST-AKT1 for 30 minutes.

Phosphorylation of ALG3 at Ser11/Ser13 has been detected in multiple independent high throughput proteomic studies (**Fig 1A**). The Ser11/Ser13 consensus motifs are conserved in mammalian species, including human, mouse and rat (**Fig. 1B**). To determine if ALG3 is phosphorylated downstream of PI3K/AKT, HA/FLAG-tagged ALG3 was expressed in MCF10A immortalized mammary epithelial cells. Cells were serum-starved, stimulated with insulin over a time course, ALG3 immunoprecipitated and phosphorylation detected with a phospho-specific antibody that recognizes the AKT consensus motif. Insulin stimulated ALG3 phosphorylation in a time-dependent manner, coincident with PI3K/AKT activation as measured by pAKT (Ser473) and the canonical substrate PRAS40 (Ser246) (**Fig. 1C**). ALG3 phosphorylation in response to insulin was blocked by GDC-0941 (pan-PI3K inhibitor), GDC-0068 (pan-AKT inhibitor), but not by rapamycin (MTORC1 inhibitor), suggesting that ALG3 phosphorylation occurs downstream of AKT, but not S6K (**Fig 1D**). Furthermore, insulin-stimulated phosphorylation of ALG3 was not observed in individual ALG3.Ser11Ala, ALG3.Ser13Ala and compound ALG3.Ser11Ala/Aer13Ala mutants, compared to wild-type ALG3 (**Fig. 1E**). In an in vitro protein kinase assay, recombinant active GST-AKT1 phosphorylated immuno-isolated recombinant wild-type ALG3, but not individual ALG3.Ser11Ala, ALG3.Ser13Ala and compound ALG3.Ser11Ala/Ser13Ala mutants (**Fig. 1F**). Collectively these data demonstrate that ALG3 is an AKT substrate downstream of PI3K, and is phosphorylated at Ser11/Ser13, in agreement with global phosphoproteomic analyses.

### ALG3 regulates protein N-glycosylation

To determine whether ALG3 functionally influences protein N-glycosylation, lectin staining was used to profile whole cell lysates in MDA-MB-468 breast cancer cells with hyperactive AKT activity due to *PTEN* inactivation. CRISPR/Cas9 pooled cell lines were generated using two independent guides for ALG3 (sgALG3_2 and sgALG3_3), and control empty vector. ALG3 knockout was confirmed by quantitative RT-PCR due to the lack of an available antibody that robustly detects endogenous AGL3 (**Fig. 2A**). Cells were grown over a period of eight days through selection, harvested and whole cell lysates profiled by lectin staining using three distinct lectins that recognize glycans inclusive of α-1,3 mannose: *Galanthus nivalis* agglutinin (GNA), Concanavalin A (ConA) and *Griffonia Simplifigfolia* Lectin I (GSL-I). GNA binds hybrid glycans, specifically terminal α-1,3 mannose residues^16^. ConA lectin recognizes α-mannose-containing cores and oligomannose-type N-glycans with a higher affinity than complex-type N-glycans, but not highly branched complex-type N-glycans nor O-glycans^16^. GSL-I binds terminal Galα and GALNAcα^16^.

**Figure 2.**
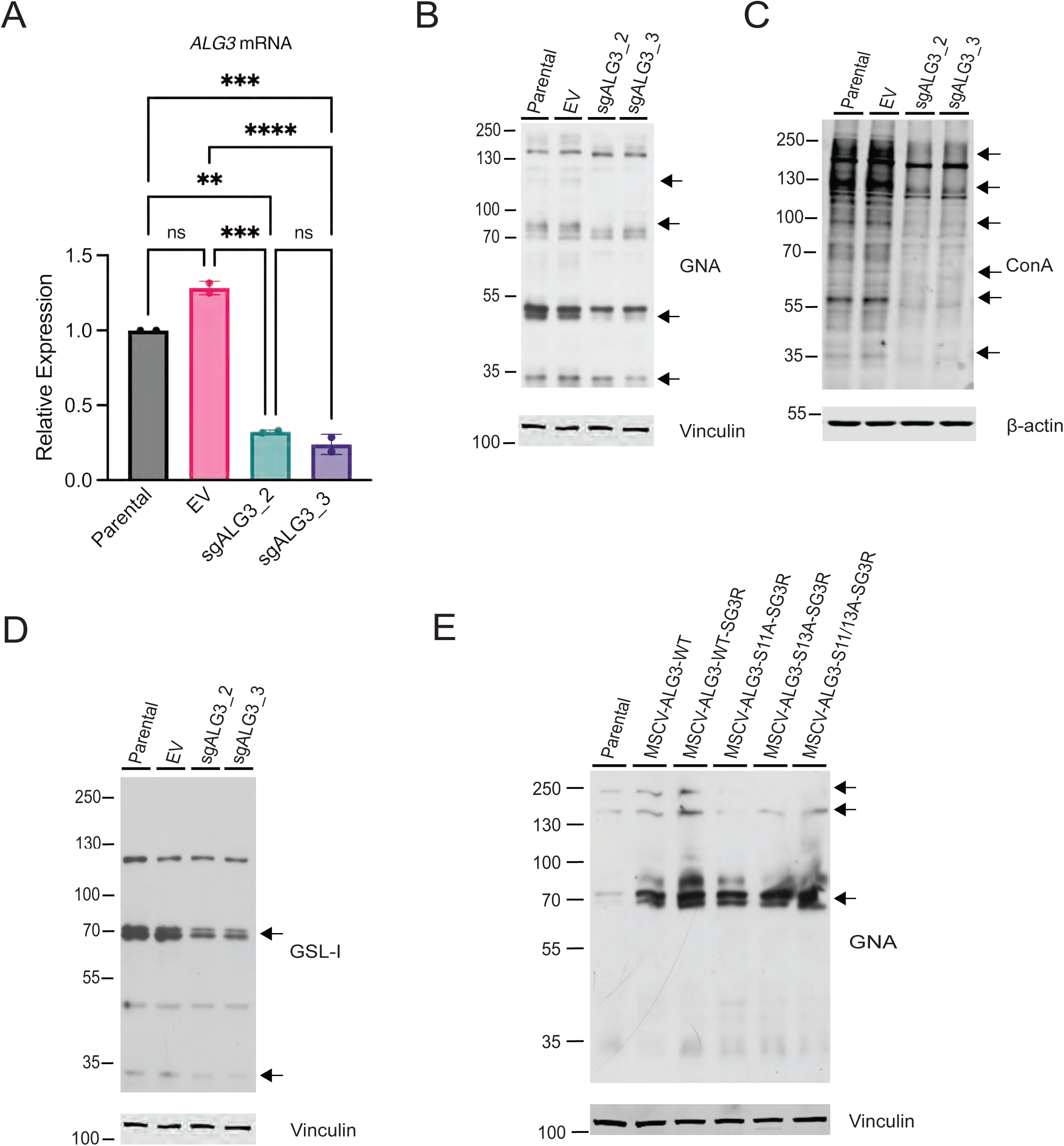
ALG3 expression levels influence lectin binding. MDA-MB-468 parental cells and cells expressing CRISPR EV or guides sgALG3_2 or sgALG3_3. (A) *ALG3* mRNA abundance was measured by qRT-PCR using two primer sets with two technical replicates and is expressed as fold change relative to parental cells. Statistical analysis was performed using two-way analysis of variance (ANOVA) with Tukey’s multiple comparisons test. sgALG3_2 and _3, **, p = 0.0019, ***, p = 0.008; and EV ***, p = 0.002, **** p < 0.001. Lectin staining was performed using (B) GNA, (C) ConA, and (D) GSL-I. (E) MCF10A parental cells and cells expressing HA-FLAG-ALG3-WT, -WT-SG3R, -S11A-SG3R, -S13A-SG3R and -S11/13A-SG3R were stained for GNA. Arrows indicate proteins with differential lectin staining.

CRISPR-Cas9 depletion of ALG3 reduced the staining pattern of all three lectins, indicating that ALG3 is functionally required for protein glycosylation using α-1,3 mannose in breast cancer cells with hyperactive PI3K signaling (**Fig. 2 B,C,D**). Specifically, decreased staining of bands migrating at 120kDa, 80kDa, 50kDa and 30kDa was observed with GNA; 200kDa, 120kDa, 100kDa, 60kDa, 57kDa and 35kDa with ConA; 70kDa and 33 kDa with GSL-1. Ectopic expression of WT-ALG3 in MCF10A cells increased the pattern of protein bands recognized by GNA lectin at 70kDa, 150kDa and 240kDa. Guide-resistant versions of WT, Ser11Ala, Ser13Ala and Ser11Ala/Ser13Ala were also expressed and similarly enhanced GNA staining of all three proteins, however the phospho-mutant alleles did not show enhanced staining of proteins at 150kDa and 240kDa (**Fig. 2E**). This suggests that phosphorylation of Ser11/Ser13 regulates the ability of ALG3 to modulate the glycome processed within the N-glycosylation pathway.

### ALG3 depletion induces ER stress and the UPR

Protein glycosylation occurs in the ER and is both necessary and sufficient for proper protein folding and ER homeostasis. Enhanced N-glycosylation supports increased protein turnover and renewal required by rapidly proliferating cells^4,14,15,36^. By contrast, impaired N-glycosylation results in accumulation of unfolded proteins, leading to ER stress and activation of the UPR, or cell death when ER stress cannot be resolved^54^.

CRISPR-Cas9 depletion of ALG3 (sgALG3_2 and sgALG3_3) led to induction of ER stress in MDA-MB-468 breast cancer cells, as determined by increased protein (**Figure 3A**) and mRNA (**Figure 3B**) of the ER stress and UPR marker 78 kDA glucose-regulated protein/binding immunoglobulin protein (*GRP78/BiP*). Moreover, a corresponding increase in the mRNA levels of the ER/UPR marker C/EBP homologous protein (*CHOP*) was also observed upon ALG3 depletion (**Figure 3B**).

**Figure 3.**
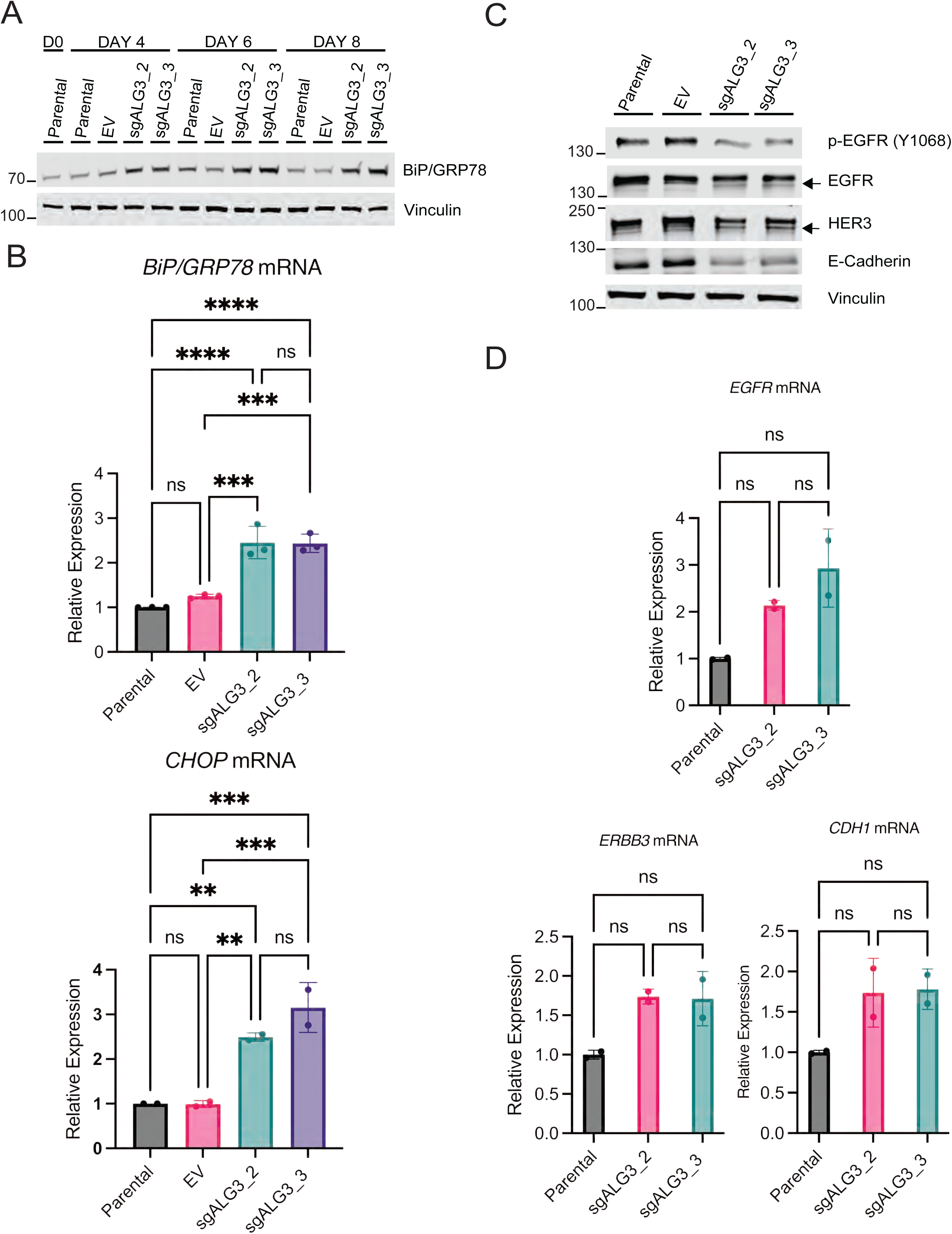
ALG3 depletion induces ER stress and attenuates receptor glycosylation. (A) whole cell lysates of MDA-MB-468 parental, EV, and sgALG3_2 and sgALG3_3 cells evaluated by immunoblotting for BiP/GRP78 and (B) by qRT-PCR for *BiP/GRP78* and *CHOP* mRNA using three primer sets with two technical replicates. mRNA levels are expressed as fold change relative to the parental line and normalized to *GAPDH* mRNA and *18S* mRNA. Statistical analysis was performed using two-way ANOVA with Tukey’s multiple comparisons test; asterisks (*) indicate significant differences in *BiP/GRP78* mRNA levels for knockout cells compared to parental (****, p < 0.0001) and EV cells (***, p = 0.0002) and indicate significant differences in *CHOP* mRNA levels for knockout cells compared to parental (**, p = 0.0033; ***, p = 0.0003) and EV cells (**, p = 0.0032; *** p = 0.0003). (C) Immunoblotting of MDA-MB-468 cells transduced with EV or sgALG3_2/3. Arrows indicate migration of deglycosylated receptors. (D) qRT-PCR analysis of mRNA abundance in MDA-MB-468 cells expressing EV or sgALG3_2/3, four days post-selection. mRNA levels are expressed as fold change relative to the parental line and normalized to *18S* mRNA. Statistical analysis was performed using one-way ANOVA with Tukey’s multiple comparisons test.

To determine the functional consequence of ALG3 depletion on proper protein folding, we evaluated a number of integral membrane proteins known to be glycosylated, including epidermal growth factor-receptor (EGFR, 11 N-glycosylation sites), human epidermal growth factor receptor 3 (HER3, 10 N-glycosylation sites), and E-cadherin (4 N-glycosylation sites). Depletion of ALG3 in MDA-MB-468 cells resulted in reduced levels of phosphorylated EGFR and E-cadherin, indicative of improper protein folding. Moreover, for both EGFR and HER3, a faster migrating band was resolved by SDS PAGE indicative of deglycosylated receptors (**Figure 3C**). The mRNA levels of *EGFR*, *ERBB3* (HER3) and *CDH1* (E-cadherin) were unaffected by ALG3 depletion (**Figure 3D**).

### ALG3 is functionally required for cell proliferation

We next determined whether ALG3 is required for cell proliferation in cells harboring hyperactivated AKT signaling. First, in wild-type MCF10A mammary epithelial cells grown in full serum, upon depletion of ALG3 via CRISPR knockout with two independent guides (sgALG3_2, sgALG3_3) cell proliferation was markedly attenuated compared to parental and control cells (**Figure 4A**). Similarly, depletion of ALG3 also significantly reduced cell proliferation in *PTEN*-deficient, AKT-hyperactivated MDA-MB-468 breast cancer cells (**Figure 4B**). Next, MDA-MB-468 parental cells and cells overexpressing WT and phosphomutant ALG3 were infected with sgALG3_3, then grown alongside their parental counterparts for seven days. For all lines, depletion of ALG3 resulted in decreased proliferation. While overexpression of WT-ALG3 resulted in a modest increase in cell proliferation relative to parental cells, overexpression of phosphomutant Ser11Ala/Ser13Ala-ALG3 resulted in decreased cell proliferation relative to parental cells, suggesting that Ser11/Ser13 phosphorylation is critical for cell growth (**Figure 4C**). Furthermore, inhibition of proliferation in MDA-MB-468 cell transduced with CRISPR-Cas9 guides sgALG3_2 and sgALG3_3 could be rescued with a wild-type ALG3 allele (ALG3-WT-SG3R) resistant to guide sgALG3_3, but not resistant to guide sgALG3_2 (**Figure 4D**). Similarly, MDA-MB-468 cells harboring an SG3 guide-resistant ALG3 ser11ala/ser13ala phosphomutant (ALG3-S11/13A-SG3R) still rescued proliferation, albeit to a lesser extent than to cells expressing guide-resistant wild-type ALG3 (in the context of vector-transduced cells compared to guide sgALG3_2; **Figure 4E**). Finally, expression of guide-resistant WT ALG3 (ALG3-WT-SG3R) in MDA-MB-468 cells transduced with guide sgALG3_3 rescued total EGFR protein levels, EGFR Y1068 phosphorylation and E-cadherin total protein, as well as the GNA staining pattern. By contrast, expression of guide-resistant phosphomutant ALG3 (ALG3-S11/13A-SG3R) failed to rescue, suggesting that ALG3 phosphorylation at Ser11/Ser13 is required for ALG3 function in glycosylation and protein folding of these glycoproteins (**Figure 4F**).

**Figure 4.**
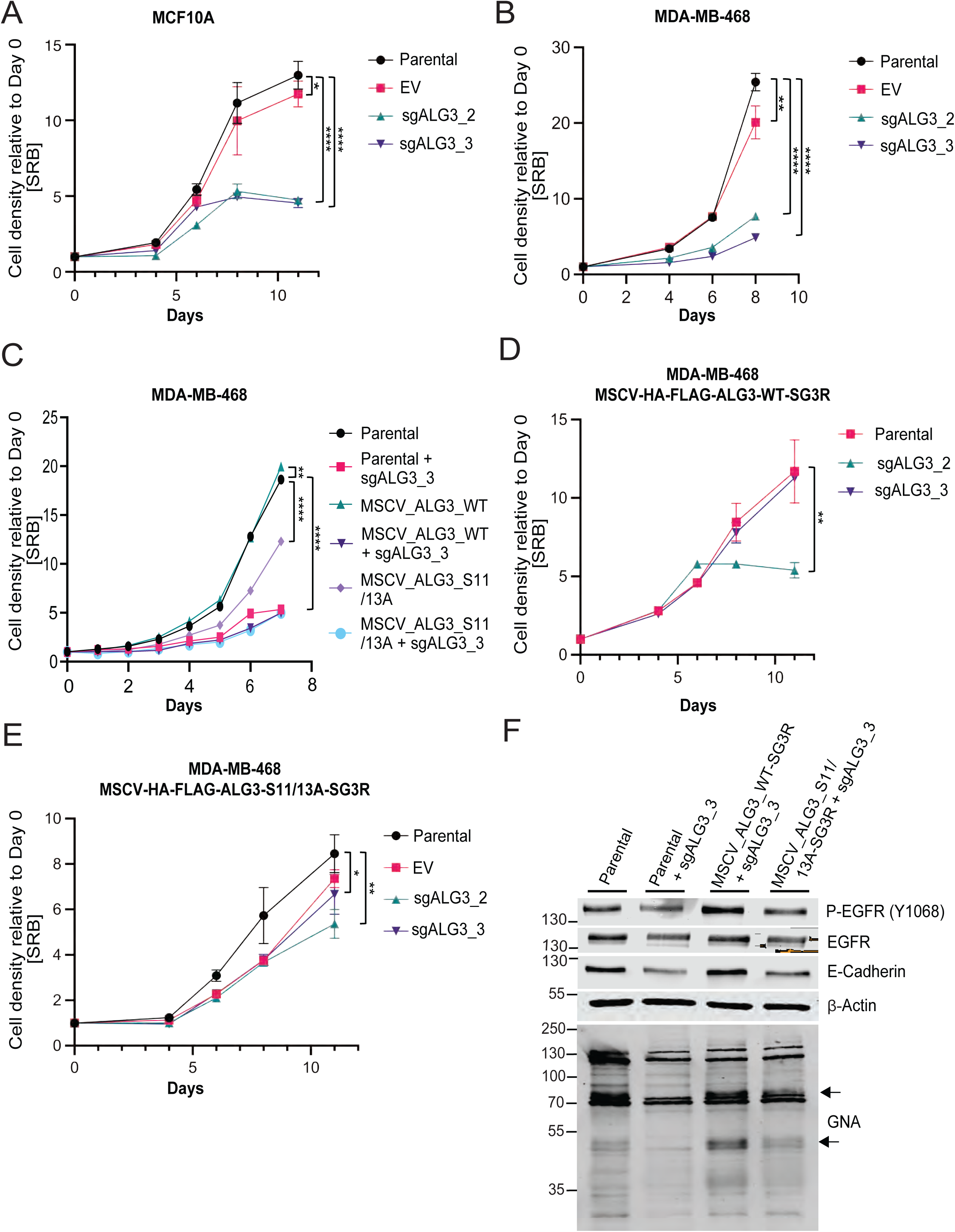
Depletion of ALG3 reduces proliferation of MCF10A and MDA-MB-468 cells. (A) SRB proliferation assay for parental cells and cells expressing EV or sgALG3_2/3 in MCF10A (*, p = 0.0381; ****, p < 0.0001) and (B) MDA-MB-468 cells (**, p = 0.0020; ****, p < 0.0001). (C) SRB assay for MDA-MB-468 parental cells and cells overexpressing HA-FLAG-ALG3-WT or HA-FLAG-ALG3-S11/13A, infected with sgALG3_3 alongside respective parental counterparts (**, p = 0.0091; ****, p <0.0001). (D) SRB for MDA-MB-468 cells expressing HA-FLAG-ALG3-WT-SG3R transduced with EV or sgALG3_2 and sgALG3_3 (**, p = 0.0012). (E) SRB for MDA-MB-468 cells expressing HA-FLAG-ALG3-S11/13A-SG3R transduced with EV or sgALG3_2 and sgALG3_3 (**, p = 0.0018; *, p = 0.0371). (F) Immunoblotting of MDA-MB-468 cells and parental, HA-FLAG-ALG3-WT-SG3R and HA-FLAG-ALG3-S11/13A-SG3R cells infected with sgALG3_3 for p-EGFR (pY1068), EGFR, E-Cadherin, β-actin, and GNA lectin. For A-D, data are expressed as mean ± SD (N = 3 technical replicates). Statistical analysis was performed using endpoint data, an ordinary one-way ANOVA and Dunnett’s multiple comparisons test.

## DISCUSSION

Both N-glycosylation and PI3K/AKT signaling are critical for normal proliferation and growth, and their deregulation contributes to tumorigenesis. Here we show that growth factor and oncogene signaling through PI3K/AKT regulates N-glycosylation to support increased protein folding needs of rapidly-proliferating cancer cells. We show that ALG3 is phosphorylated at Ser11 and Ser13 by AKT in growth factor-stimulated cells, and that phosphorylation of ALG3 is required for proper glycan addition and protein folding of cell surface receptors. Although AKT typically inhibits substrates upon phosphorylation, it also upregulates anabolic pathways such as lipid, protein and nucleotide metabolism to satisfy the needs of rapidly proliferating cells^6,55^. Our data are consistent with a model in which phosphorylation of ALG3 by AKT is an activating event, accelerating N-glycan production to meet the demands of oncogenic signaling and proliferation. This is consistent with the finding that inhibition of ALG3 leads to induction of the unfolded protein response, endoplasmic reticulum stress, and impaired cell proliferation in PI3K/AKT hyperactive cancer cells. Impaired phosphorylation of ALG3 using Ser11/Ser13 phosphomutants only modestly affected cell proliferation, at least under our experimental conditions, indicating that other modifications may contribute to N-glycosylation and activity. By contrast, impaired ALG3 Ser11/Ser13 phosphorylation did attenuate protein glycosylation as measured by lectin staining and reduced expression of deglycosylated receptors such as EGFR and HER3.

Prior studies have suggested a functional role for ALG3 in malignancy. In *Drosophila*, ALG3 loss disrupts N-glycosylation of the tumor necrosis factor receptor, elevated c-Jun N-terminal kinase, suppression of the Hippo pathway and increased cell proliferation^56,57^. In humans, *ALG3* gene amplification, which has been associated with poor outcomes in breast cancer, has also been observed in lung, ovarian, neuroendocrine prostate, cervical, bladder, endometrial, head and neck, and esophageal cancers^51,58–60^. Interestingly, the *ALG3* gene resides on the same 3q26-27 amplicon as *PIK3CA*, which encodes the p110α catalytic subunit of class I PI3K. Consequently, *ALG3* and *PIK3CA* are frequently coamplified in many human carcinomas^58,59^. Moreover, while ALG3 overexpression promotes cancer progression, loss-of-function *ALG3* mutations result in severe clinical manifestations in congenital disorders of N-glycosylation (ALG3-CDG). These patients harbor compound heterozygous *ALG3* mutations, but homozygous mutations are rarely found suggesting these may be lethal during embryonic development^43^. ALG3-CDG patients display developmental and intellectual disabilities, muscular hypotonia, cerebral malformations, neurological features such as epileptic seizures, cardiac defects, facial and body dysmorphism, and metabolic dysfunction including refractory hypoglycemia^43,45^. Molecularly, ALG3-CDG patients have accumulated truncated Man5-GlCNAc2 structures, expected to be found in cells lacking functional ALG3^47^. Taken together, these findings underscore a critical balance in ALG3 expression and function.

Collectively, we propose a model in which in addition to anabolic protein, nucleotide and lipid metabolism, the PI3K/AKT pathway promotes N-glycosylation through ALG3 to facilitate proper folding of proteins destined for secretion and the cell surface in rapidly proliferating cells, including cancer. Identifying the precise identity of the glycome and the proteoglycome that is regulated by PI3K/AKT/ALG3 using, for example, peptidoglycomics, is a future goal and would uncover a new layer on the role of AKT signaling in cancer. These findings also raise the question as to whether additional glycosyltransferases are also functionally modified by post-translational modifications. In this context, in silico data mining reveals that, for example, α-1,3/1,6-mannosyltransferase (ALG2) also harbors an AKT substrate consensus that has been mapped by high-throughput phospho-proteomic studies (Ser256)^61^. Similarly, a link between MEK-ERK signaling and N-glycosylation is suggested by the fact that the dolichyl-phosphate mannosyltransferase subunit 1 (DPM1) harbors multiple putative proline-directed kinase (e.g. ERK, CDK) phosphorylation sites, again mapped by functional phosphoproteomics^62,63^. DPM1 catalyzes the first committed-step step in the N-glycosylation pathway, and is also amplified in ovarian and colon cancer^16,64,65^. These represent just two examples of glysosyltransferases that are likely modified by Ser/Thr phosphorylation in cells and tissues, raising the possibility that functional modulation of the glycome and proteglycome lies downstream of oncogene-driven signaling pathways such as PI3K/AKT/MTORC1 and RAS/MEK/ERK.

Understanding the regulation of N-glycosylation by signaling pathways such as PI3K/AKT may also afford new therapeutic interventions in diseases such as cancer and congenital disorders of N-glycosylation. Deregulated glycosylation has been associated with a failure to thrive in infants, impaired neurological development and infantile-onset symptomatic epilepsy^21,66^. Uncovering how signaling pathways regulate glycosylation and how aberrant signaling can alter glycosylation may provide new approaches for the combination strategies targeting both protein kinases and glycosylation. As but one example, drugs that inhibit oligosaccharyltransferase, the enzyme that transfers oligosaccharides to proteins to facilitate folding, have shown promise in non-small-cell lung cancer by selectively blocking EGFR^67^. Targeting ALG3 may similarly provide selective inhibition of N-glycosylation, potentially improving therapeutic efficacy. Collectively, these findings advance our understanding of the regulation of N-glycosylation by growth factor and oncogene-driven signaling and its role in normal cellular physiology as well as cancer.

## Abbreviations

MTORC1: mechanistic target of rapamycin
PI3K: phosphoinositide 3-kinase
PTEN: phosphatase and tensin homolog
INPP4B: inositol polyphosphate 4-phosphatase type II
ER: endoplasmic reticulum
ALG3: asparagine-linked glycosylation 3 homolog
CDG: congenital disorders of N-linked glycosylation
PTM: post-translational modification
UPR: unfolded protein response
EV: empty vector
GNA: *Galanthus nivalis agglutinin*
ConA: *Concanavalin A*
GSL-I: *Griffonia (Bandeiraea) Simplifigfolia* Lectin I
GRP78/BiP: 78 kDA glucose-regulated protein
CHOP: C/EBP homologous protein
EGFR: epidermal growth factor-receptor
HER3: human epidermal growth factor receptor 3
ALG2: α-1,3/1,6-mannosyltransferase
DPM1: dolichyl-phosphate mannosyltransferase subunit 1
ANOVA: analysis of variance

## ACKNOWLEDGMENTS

This work was supported by the following grants: CA253097 (A.T.), F31CA250094 (A.J.N.). We thank members of the Toker lab for constructive comments and Dr. Richard Cummings (BIDMC), Dr. Davie Van Vactor (HMS), Dr. Dohoon Kim (UMass), Dr. Steve Elledge (HMS), Dr. Kevin Haigis (DFCI) and Dr. Andrew Aguirre (DFCI) for helpful suggestions and project advice.

## AUTHOR CONTRIBUTIONS

A.T., E.C.L., L.G., and A.J.N. conceptualization; A.J.N., E.C.L., and L.G. methodology; A.J.N. formal analysis; A.J.N. investigation; A.J.N. writing original draft; A.J.N. and A.T. writing, review and editing; A.T. supervision; A.T. resources; A.J.N. visualization; A.T. project administration; A.T. funding acquisition.

## DATA AVAILABILITY

Source data and annotated analysis workflows are available on the following OSF project website: https://osf.io/v4tzs/.

## EXPERIMENTAL PROCEDURES

Detailed experimental procedures, including protocols for drug treatment experiments, immunoblotting, immunoprecipitation, CRISPR knockouts, site-directed metagenesis, overexpression, lectin blotting, in vitro kinase assay, immunofluorescence, and qRT-PCR, can be found in the supporting information.

## CONTACT FOR REAGENT AND RESOURCE SHARING

Further information and requests for resources and reagents should be directed to and will be fulfilled by the Lead Contact, Alex Toker. Requested compounds will be provided following completion of an MTA.

## EXPERIMENTAL MODELS AND SUBJECT DETAILS

### CELL CULTURE

MCF10A cells were cultured in standard MCF10A growth medium without antibiotics [DMEM/F12 medium (Wisent Bioproducts, 319-075-CL)], 5% horse serum (Gemini Bio, 100508), 10 µg ml^-1^ insulin (Thermo Fisher Scientific/Gibco, A1138211), 0.5 mg ml^-1^ hydrocortisone (Sigma-Aldrich,H4001), 20 ng ml^-1^ EGF (R&D Systems, 236-EG-01M) and 100 ng ml^-1^ cholera toxin (List Biological Laboratories, 100B). All MDA-MB-468-derived cell lines were cultured in RPMI media (Wisent Bioproducts) supplemented with 10% heat-inactivated fetal bovine serum (Thermo Fisher Scientific) without antibiotics. All cell lines were cultured at 37°C in the presence of 5% CO_2_. Cells were passaged for no more than 4 months and routinely assayed for mycoplasma contamination.

### INHIBITORS AND GROWTH FACTORS

For signaling experiments, serum/growth-factor-deprived cells (20 hours) were pre-treated for 15 minutes with DMSO (Thermo Fisher Scientific, BP231-100) or small-molecule inhibitors, then treated with insulin (Thermo Fisher Scientific/Gibco, A1138SII [dissolved in 0.1N HCl then diluted in water to final concentration]) for the indicated time period. Stocks were prepared under sterile conditions, aliquoted and stored at –80C. The small-molecule kinase inhibitors used included the PI3K catalytic inhibitor GDC-0941 (Cayman Chemical, 11600), the AKT catalytic inhibitor GDC-0068 (Selleck Chemicals, S2808) which were each dissolved in DMSO at 10mM. The mTORC1 inhibitor Rapamycin (Cayman Chemical, 13346) was dissolved in DMSO at 100uM. All inhibitors were aliquoted under sterile conditions, aliquoted and stored at –80C.

### ANTIBODIES AND LECTINS

The following antibodies were used: P-AKT substrate (RXXS*/T*) (9614S, 1:1000), AKT (pan) (4691S, 1:1000), P-AKT (S473) (4060L, 1:1000), P-PRAS40 (T246) (2997S, 1:1000), PRAS40 (tot) (2691S, 1:1000), vinculin (12901S, 1:1000), p-p70 S6 Kinase (T389) (9234S, 1:1000), p70 S6 Kinase (2708S, 1:1000), PERK (3192S, 1:1000), BiP/GRP78 (3177S, 1:1000), P-EGFR (Y1068) (3777S, 1:1000), EGF Receptor (4267S, 1:1000), HER3 (12708S, 1:1000), E-Cadherin (3195S, 1:1000), GST (2625S, 1:1000), and HA-Tag (2367S, 1:1000) were purchased from Cell Signaling Technology. ALG3-1 (Q92685_1, 1:300) was custom made by GenScript. B-Actin (067M4856V, 1:1000) was purchased from Sigma. The following lectins were used: Galanthus Nivalis Lectin (GNL/GNA), Biotinylated (B-1245-2, 1:750), Concanavalin A (Con A), Biotinylated (B-1005-5, 1:1500), and Griffonia (Bandeiraea) Simplicifolia lectin I (GSL-I), (BK-2100, 1:750) were all purchased from Vector Laboratories.

### QUANTIFICATION AND STATISTICAL ANALYSIS

Statistical analyses were performed with PRISM 10 graphing software (GraphPad). N=3 technical and biological replicates were used and each data point represents the mean +/− SEM. Statistical analyses for SRB growth curves were performed by completing a one-way ANOVA at the latest time point.

