## Supporting methods for "The glycosyltransferase ALG3 is an AKT substrate that regulates protein N-glycosylation"

**Supporting Information – Full Methods Description**

**Cell Culture.**

MCF10A cells were cultured in standard MCF10A growth medium without antibiotics (DMEM/F12 medium (Wisent Bioproducts, 319-075-CL), 5% horse serum (Gemini Bio, 100508), 10 µg $ml^{-1}$ insulin (Thermo Fisher Scientific/Gibco, A1138211), 0.5 mg $ml^{-1}$ hydrocortisone (Sigma-Aldrich,H4001), 20 ng $ml^{-1}$ EGF (R&D Systems, 236-EG-01M) and 100 ng $ml^{-1}$ cholera toxin (List Biological Laboratories, 100B). All MDA-MB-468-derived cell lines were cultured in RPMI media (Wisent Bioproducts) supplemented with 10% heat-inactivated fetal bovine serum (Thermo Fisher Scientific) without antibiotics. All cell lines were cultures at 37°C in the presence of 5% $CO_2$. Cells were passaged for no more than 4 months and routinely assayed for mycoplasma contamination.

**Inhibitors and Growth Factors.**

For signaling experiments, serum/growth-factor-deprived cells (20 hours) were pre-treated for 15 minutes with DMSO (Thermo Fisher Scientific, BP231-100) or small-molecule inhibitors, then treated with insulin (Thermo Fisher Scientific/Gibco, A1138SII, final concentration 100nM, stock concentration 100 uM [dissolved in 0.1N HCl then diluted in water to final concentration]) for the indicated time period. Stocks were prepared under sterile conditions, aliquoted and stored at –80C. The small-molecule kinase inhibitors used included the PI3K catalytic inhibitor GDC-0941 (Cayman Chemical, 11600), the AKT catalytic inhibitor GDC-0068 (Selleck Chemicals, S2808) which were each dissolved in DMSO at 10mM. The mTORC1 inhibitor Rapamycin (Cayman Chemical, 13346) was dissolved in DMSO at 100uM . All inhibitors were aliquoted under sterile conditions, aliquoted and stored at –80C. Aliquots were freeze-thawed no more than four times.

**Antibodies.**

P-AKT substrate (RXXS*/T*) (9614S, 1:1000), AKT (pan) (4691S, 1:1000), P-AKT (S473) (4060L, 1:1000), P-PRAS40 (T246) (2997S, 1:1000), PRAS40 (tot) (2691S, 1:1000), vinculin (12901S, 1:1000), p-p70 S6 Kinase (T389) (9234S, 1:1000), p70 S6 Kinase (2708S, 1:1000), PERK (3192S, 1:1000), BiP/GRP78 (3177S, 1:1000), P-EGFR (Y1068) (3777S, 1:1000), EGF Receptor (4267S, 1:1000), PD-L1

(13684S, 1:1000), HER3 (12708S, 1:1000), E-Cadherin (3195S, 1:1000), GST (2625S, 1:1000), and HA-Tag (2367S, 1:1000) were purchased from Cell Signaling Technology. ALG3-1 (Q92685_1, 1:300) was custom made by GenScript. B-Actin (067M4856V, 1:1000) was purchased from Sigma.

**Lectins.**

Galanthus Nivalis Lectin (GNL/GNA), Biotinylated (B-1245-2, 1:750), Concanavalin A (Con A), Biotinylated (B-1005-5, 1:1500), Phaseolus Vulgaris Erythroagglutinin (PHA-E) , (B-1125-2, 1:750), Sambucus Nigra Lectin (SNA, EBL), Biotinylated (B-1305-2, 1:750), Griffonia (Bandeiraea) Simplicifolia lectin I (GSL-I), (BK-2100, 1:750), Ricinus Communis Agglutinin (RCA-I), Biotinylated (BK-1000, 1:750), and Ulex Europaeus Agglutinin (UEA-I), Biotinylated (BK-1000, 1:750) were all purchased from Vector Laboratories.

**Drug Treatment Experiments.**

For signaling experiments cells were plated in 15 cm plates. Two days later, cells are serum/growth-factor-deprived (20 hours). Cells are then pre-treated for 15 minutes with DMSO  or small-molecule inhibitors (GDC0941, GDC0068, or Rapamycin), then treated with insulin for the indicated time period. Cells were then harvested as described in 'Immunoprecipitation analysis of Endogenous ALG3'.

**Immunoblots.**

Cells were washed with 1X PBS (Boston Bioproducts) and then either flash frozen and stored at –80C until lysis, or lysed fresh, using radioimmunoprecipitation buffer (RIPA) (1% Nonidet P-40, 0.5% sodium deoxycholate, 0.1% SDS, 150 mM NaCl, 50 mM Tris-HCl, pH 7.5, protease inhibitor mixture, 50 nM calyculin A, 1mM sodium pyrophosphate, and 20mM sodium fluoride) for 15 minutes. Cell extracts were precleared by centrifugation at 14,000 rpm for 10 min at 4C, and then cleared by a second centrifugation at 14,000 rpm for 10 min at 4C. The Bio-Rad DC protein assay was used to measure the protein concentration, and then sample concentration was normalized using RIPA and SDS sample buffer. All samples in which ALG3-1 is detected are not boiled or frozen and loaded fresh. All other samples are boiled for 5 min at 95C. Lysates were resolved on acrylamide gels by SDS-polyacrylamide gel electrophoresis and electrophoretically transferred to nitrocellulose membrane (BioRad) at 300 mA for 90 minutes at 4C in Tris-glycine transfer buffer (Boston BioProducts). Membranes were blocked in TBS buffer (10 mmol/L Tris-HCl, pH 8, 150 mmol/L NaCl; Boston Bioproducts) containing 5% (w/v) nonfat dry milk (Andwin Scientific). Membranes were incubated with specific primary antibodies diluted in 5% (w/v) fatty-acid-free bovine serum albumin (BSA) (Boston BioProducts, P-753) in TBS-T with 0.01% (w/v) sodium azide (Thermo Fisher Scientific) and rocking at 4C overnight. Membranes were washed in TBST at room temperature three times for 10 minutes and if developed in the dark room were incubated in horseradish peroxidase (HRP)-conjugated secondary antibody diluted in 5% (w/v) milk in TBS-T for 1 hour, with rocking at room temperature. The membrane was washed again with TBS-T then incubated in Clarity Western ECL Substrate (Bio-Rad, 1705061) or Clarity MAX Western ECL Substrate (Bio-Rad, 1705062), then developed. If the membrane was not incubated with HRP secondary, it was incubated in fluorophore-conjugated secondary antibodies (LI-COR Biosciences) diluted in 5% (w/v) milk in TBS-T for 1 hour, with rocking at room temperature. The membrane was washed again with TBS-T then the chemiluminescence signal was imaged with a Li-Cor Odyssey CLx Imaging System (LI-COR Biosciences).

**Immunoprecipitation analysis of endogenous ALG3.**

All steps were carried out on wet ice or at 4C. Cells were lysed from 15 cm dishes in 100 ul of RIPA buffer (1% Nonidet P-40, 0.5% sodium deoxycholate, 0.1% SDS, 150 mM NaCl, 50 mM Tris-HCl, pH 7.5, protease inhibitor mixture, 50 nM calyculin A, 1mM sodium pyrophosphate, and 20mM sodium fluoride) on ice for 15 min. Lysates were centrifuged at 14,000 rpm for preclearing for 10 min at 4C, the supernatants transferred to a new tube, then centrifuged a second time at 14,000 rpm for 10 minutes at 4C, and the cleared supernatants were transferred to a new tube. The Bio-Rad DC protein assay was used to assess protein concentration, and sample concentrations were normalized using RIPA and SDS sample buffer. 1.4 mg of lysate was incubated for 2 hours with immunoprecipitation antibody, rocking at 4C. Immunoprecipitation antibodies were bound to Protein A/G agarose beads that had been washed three times with PBS, then resuspended in a 1:1 slurry in RIPA. Additional lysate was saved for immunoblotting the whole cell lysate. Then, 20 ul of resuspended beads were added to each immunoprecipitation sample for an additional 2 hours with rocking at 4C. Three washes were performed in RIPA lysis buffer and beads were centrifuged at 2,500g, followed by removal of all of the supernatant. One wash was performed using PBS. The samples were then resuspended in 1X sample buffer and heated to 65C for 20 minutes to release the protein from the beads, then loaded on acrylamide gels by SDS-polyacrylamide gel electrophoresis. Samples are not boiled or frozen.

**Immunoprecipitation analysis of MSCV-HA-FLAG-ALG3.**

Samples were lysed as described in the 'Immunoprecipitation analysis of endogenous ALG3' section. Anti-flag M2 agarose beads (Millipore-Sigma, A2220) were added to lysates and then samples were rocked for 2 hours at 4C to immunopurify HA-FLAG-ALG3. The beads were washed three times in full RIPA lysis buffer, and one time with 1X PNS at 4C; beads were centrifuged at 14,000 rpm, followed by removal of all of the supernatant. Excess buffer was removed from above the settled beads before resuspended in 1X sample buffer and heated to 65C for 20 minutes for release of protein from the beads then loaded on acrylamide gels by SDS-polyacrylamide gel electrophoresis. Samples are not boiled or frozen. After transfer, gels were Coomassie stained, and then incubated for 1 hour with specific primary antibodies diluted in 5% (w/v) fatty-acid-free bovine serum albumin (BSA) (Boston BioProducts, P-753) in TBS-T with 0.01% (w/v) sodium azide (Thermo Fisher Scientific) and rocking at 4C overnight. Membranes were washed in TBST at room temperature three times for 10 minutes and if developed in the dark room were incubated in horseradish peroxidase (HRP)-conjugated secondary antibody diluted in 5% (w/v) milk in TBS-T for 1 hour, with rocking at room temperature. The membrane was washed again with TBS-T then incubated in Clarity Western ECL Substrate (Bio-Rad, 1705061) or Clarity MAX Western ECL Substrate (Bio-Rad, 1705062) then developed in the dark room.

**Growth Rate Assay.**

Cell lines were plated at densities ranging from 5,000-25,000 cells per well in 24-well plates and allowed 24 hours to adhere before a day zero reading is taken. Population size as cell confluency was assayed using sulforhodamine B staining, as described in (Vichai V., and Kirtikara K. (2006) Sulforhodamine B colorimetric assay for cytotoxicity screening. Nat. Protoc. 1, 1112–1116 10.1038/nprot.2006.179). Cells were fixed in 8.3% final concentration of tricarboxylic acid for at least 1 hour at 4C at the indicated day. Cells were then washed three times with DI water. Cells were then stained for at least 30 min with 0.5% sulforhodamine B in 1% acetic acid. Cells were then washed three times with 1% acetic acid and then the sulforhodamine B dye was solubilized in 10mM Tris pH 10.5. Absorbance was read at 510 nm using the BioTek Synergy LX multi-mode plate reader. Relative growth was determined as (growth at each day/growth at day 0).

**Lentivirus production.**

293T cells were plated in 10cm dishes at $9x10^6$ cells/plate then transiently transfected with 6.3 ug of the plasmid of interest to generate lentivirus. 11.1 ug of the psPAX2 packaging plasmid (Addgene plasmid, 12260, created by the laboratory of D. Trono), 0.6 ug of pCMV-VSV-G envelope protein plasmid (Addgene plasmid 8454, created by the laboratory of B. Weinberg) and 8.6 ul of polyethylenamine (Millipore-Sigma, 408727) per ug of DNA transfected. After 48 hours, media is collected, passed through a 0.45 um filter (Thermo Fisher Scientific, 09-740-106), aliquoted to 5 mL aliquots, and stored at –80C and only thawed for use once, never re-frozen.

**Retrovirus production.**

293T cells were plated in 10 cm dishes at $9x10^6$ cells/plate then transiently transfected with 2.5 ug of the plasmid of interest to generate retrovirus. 1.5 ug of the pCL-Eco packaging plasmid (Addgene plasmid, 12260, created by the laboratory of D. Trono), 1 ug of pCMV-VSV-G envelope protein plasmid (Addgene plasmid 8454, created by the laboratory of B. Weinberg) and 21.6 ul of polyethylenamine (Millipore-Sigma, 408727) per ug of DNA transfected. After 48 hours, media is collected, passed through a 0.45 um filter (Thermo Fisher Scientific, 09-740-106), aliquoted to 5 mL aliquots, and stored at –80C and only thawed for use once, never re-frozen.

**ALG3 CRISPR-Cas9 Knockouts.**

The pLentiCRISPRv2-Blast vector was used, which would constitutively express the sgRNA and the blasticidin resistance marker (Addgene plasmid, 83480 created by the laboratory of Mohan Babu),

containing two separate single guide RNA (sgRNA) sequences targeting exon 5 (sgALG3_2) and exon 4 (sgALG3_3).The sgRNAs were designed using Benchling. Guide oligos were synthesized by Integrated DNA Technologies and inserted into the vector as previously described (Genome-scale CRISPR-Cas9 knockout screening in human cells Science 2014). The plasmids were transformed into NEB Stable competent high-efficiency Escherichia coli (New England Biolabs, C3040H) then sequenced. Lentivirus was produced as described in 'Lentivirus production'. Cells were infected with lentivirus for 48 hours, then virus was removed and cells were selected with blasticidin (Invivogen, ant-bl-1) for 5-7 days until uninfected cells were completely dead from treatment. SgRNA sequences:
CACACAGTTTGGCTTCCGTG (ALG3_2), GCGGCTCTTCAATGACCCAG (ALG3_3)

**Site-directed mutagenesis of phosphorylation sites.**

The site-directed mutagenesis of the phosphorylation sites on ALG3 were performed using self-designed primers with the support of Benchling and using the QuickChange II XL Site-Directed Mutagenesis Kit (Agilent , 200521).

**MSCV-N-Flag-HA-IRES-PURO-ALG3 expression constructs.**

The full-length human ALG3 sequence with a stop codon at the end was inserted into N-terminally HA (YPYDVPDYA) and Flag-tagged (DYKDDDDK) MSCV construct. An guide-resistant version of the MSCV-N-Flag-HA-IRES-PURO_ALG3 construct was also generated by site-directed mutagenesis to be resistant to guide sgALG3_3. Phosphomutant versions of ALG3 (Ser11Ala, Ser13Ala, Ser11Ala/Ser13Ala) were also inserted into the N-terminally HA and Flag-tagged MSCV construct and additionally, sgALG3_3-resistant phosphomutant lines were generated. Empty MSCV-N-Flag-HA-IRES-PURO was used as a control for transfections and experiments. These expression constructs were transfected using HEK293T cells to produce retrovirus that was used to infect cells (see 'Retrovirus production').

**Generation of cell lines with stable expression.**

Cells were plated in 10 cm plates at $1x6^6$ cells/plate. Lentivirus/retrovirus was added with 5 ug $ml^{-1}$ polybrene (Sigma-Aldrich, 107689) to infect cells the next day. Lentivirus/retrovirus was left on the cells for 48 hours, then removed and cells were split into selection media containing 1 ug $ml^{-1}$ puromycin (MSCV) or 5-7.5 ug $ml^{-1}$ Blasticidin (pLentiCRISPR). Cells were selected until uninfected cells containing the same dosage of puromycin/blasticidin were completely dead.

**Lectin Blotting Analysis.**

Glycoprotein samples were prepared as described in 'Immunoblots' identically through the transfer step. After transfer, membranes were blocked in 5% (w/v) fatty-acid-free bovine serum albumin (BSA) (Boston BioProducts, P-753) in TBS (ThermoFisher Scientific) overnight. The blocking buffer was removed, and membranes were then incubated with biotinylated lectin diluted in 5% w/v BSA for 30 minutes rocking at room temperature. The lectin was removed, and the membranes were washed 3 times, for 10 minutes each with TBS-T (ThermoFisher Scientific) rocking at room temperature. Washing buffer was removed and Streptavidin-HRP or Streptavidin-DyLight-800 diluted in 5% w/v BSA in TBS for 30 minutes rocking at room temperature. The membranes were then washed 3 times for 10 minutes each with TBS-T rocking at room temperature. If Streptavidin-HRP was used, membranes were then incubated with Clarity Western ECL Substrate (Bio-Rad, 1705061) or Clarity MAX Western ECL Substrate (Bio-Rad, 1705062), then developed in the dark room. If Streptavidin-DyLight-800 was used, membranes were then imaged with a Li-Cor Odyssey CLx Imaging System (LI-COR Biosciences).

***In Vitro* Kinase Assays.**

MCF10A cells were plated at $5x10^5$ cells/plate in 10A plates, The next day media was removed, cells were washed with serum/growth-factor-free media, then serum starved for 20 hours. Serum-free media was then replenished for 2 hours then cells were lysed on ice and rocked for 10 minutes at 4C. Cell extracts were precleared by centrifugation at 14,000 rpm for 10 min at 4C, and then cleared by a second centrifugation at 14,000 rpm for 10 min at 4C. The Bio-Rad DC protein assay was used to measure the protein concentration, and then 2 mg of protein was immunoprecipitated, and the rest of the cell lysate saved for whole cell lysate western blotting. 30 ul of GLAF beads were added and samples were rotated for 2 hours at 4C. Samples were spin at 8K xg at 4C for 1 minute, and then supernatant was removed. Samples were washed 4 times: 1 ml NETN buffer was added, the tube was inverted 5 times and incubated on ice for 2 min, then spun at 8K xg at 4C for 1 min and supernatant removed. Tubes were washed 1 time with 1X PBS. A mastermix of reaction mix minus enzyme was prepared for each sample using water, Kinase assay buffer (10X) (cell signaling #9802; -20°C box name kinase buffer): with freshly added DTT (final 2 mM), ATP (10 mM) (Cell signaling #9804, or home-made), and Akt1, active (Sigma SRP5001). 45 ul of reaction mix minus enzyme was added to each tube with 5 ul of GST-AKT1, tubes were mixed gently by flicking, then incubated for 1 hour at 30C. Beads were flicked every 10 minutes during this hour. Reaction was stopped by adding 12 ul 6X SDS loading buffer and samples were boiled at 65C for 20 minutes and then spun at 8K x g for 30 seconds, then loaded (30 or 35 ul) on a gel for electrophoresis and subsequent western blotting.

**Immunofluorescence.**

Cells were plated at 20,000-50,000 cells per cell in 96-well plates. Cells are washed 1X with 1X PBS then 100 ul of 4% paraformaldehyde (PFA) (Buffered pH 6.9) (Millipore-Sigma, 1004965000) is added per well to the plate for 15 min at room temperature. The PFA is removed and three 5 minute washes are performed with 1X PBS. If permeabilization is desired, cells are then permeabilized with 0.2% Triton X (Fisher, BP151-100) for 10-15 minutes at room temperature. Cells are then washed with PBS 3X for 5 minutes each rocking at room temperature. All cells, with or without permeabilization, are then blocked with 0.5% BSA for 1 hour at room temperature rocking. Cells are then incubated with primary antibody overnight rocking at 4C, with 50 ul of primary at 1:50-1:100 dilutions. Cells are then washed 3 times with PBS at room temperature rocking. Cells are then incubated with secondary antibody on the rocker for 1 hour at room temperature covered (from light). Goat-anti-rabbit-568 was used at 1:1000 and DAPI was used at 1:5000. Cells are then washed 3 times for 5 minutes each rocking at room temperature. Plates were then imaged using the Keyence Microscope.

**Quantitative real-time PCR.**

Total RNA was isolated with the NucleoSpin RNA Plus (MACHEREY-NAGEL) according to the manufacturer’s protocol. Reverse transcription was performed using the SuperScript III First-Strand Synthesis System (ThermoFisher Scientific) using random hexamers as the primer. A CFX384 Touch

Real-Time PCR Detection System (BioRad) was used to perform the quantitative real-time PCR (qRT-PCR). 18S rRNA was used as the reference gene when in calculations to quantify mRNA expression. ALG3 primers were designed using Benchling. BiP/GRP78[330,331], CHOP[330,331], HER3[331], PD-L1[331,332], EGFR[331,333], and E-Cadherin[331,334] primers were published.

Primer Sequences (F=Forward, R=Reverse):

| | |
|---|---|
| ALG3-1-F: 5'GGGACTCTGCAAGCAATGGC3' | ALG3-1-R: 5'ACCCTGTGAATGACCCAGAAGG3' |
| ALG3-2-F: 5'GGGACTCTGCAAGCAATGGC3' | ALG3-2-R: 5'CCACCCTGTGAATGACCCAGAA3' |
| ALG3-3-F: 5'CCTACATGGCCGAGGTAGAAGG3' | ALG3-3-R: 5'ACAAGTGGTCCGGTGTCACC3' |
| ALG3-4-F: 5'TGGGGTTGTACTATGCCACCAG3' | ALG3-4-R: 5'GAAGACAAGCAGCAAGGTAGCC3' |
| ALG3-5-F: 5'CGTGTCCACTCCATCTTTGTGC3' | ALG3-5-R: 5'GCCAGCAGGAGGTTGATACTGA3' |
| BiP-1-F: 5'CTGTCCAGGCTGGTGTGCTCT3'. | BiP-1-R: 5'CTTGGTAGGCACCACTGTGTTC3' |
| BiP-2-F: 5'TCTGGTACTGCTTGATGTTTGTC3'. | BiP-2-R: 5'GTCCGCATCCTGGTGGCTTTCC3' |
| BiP-3-F: 5'TGGGTCGACTCGAATTCCAAAG3' | BiP-3-R: 5' GTCAGGCGATTCTGGTCATTGG |
| CHOP-1-F: 5'GGTATGAGGACCTGCAAGAGGT3' | CHOP-1-R: 5' CTTGTGACCTCTGCTGGTTCTG3' |
| CHOP-2-F: 5'TGCCTTTCACCTTGGAGAC3'' | CHOP-2-R: 5' CGTTTCCTGGGGATGAGATA3' |
| HER3-F: 5'CTATGAGGCGATACTTGGAACGG3' | HER3-R: 5'GCACAGTTCCAAAGACACCCGA3' |
| PD-L1-F: 5'TGCCGACTACAAGCGAATTACTG3' | PD-L1-R: 5'CTGCTTGTCCAGATGACTTCGG3' |
| EGFR-F: 5'AACACCCTGGTCTGGAAGTACG3' | EGFR-R: 5'TCGTTGGACAGCCTTCAAGACC3' |
| E-Cad-F: 5'GCCTCCTGAAAAGAGAGTGGAAG3' | E-Cad-R: 5'TGGCAGTGTCTCTCCAAATCCG3' |
| 18S-F: 5'-CTTAGAGGGACAAGTGGCG-3' | 18S-R: 5'-ACGCTGAGCCAGTCAGTGTA-3' |

**General Chemistry Methods.**

Reagents and solvents were purchased from commercial suppliers and were used without further purification unless otherwise noted.

**Quantification and Statistical Analysis - Proliferation Assay.**

Statistical analyses were performed with PRISM 10 graphing software (GraphPad). N=3 technical and biological replicates were used and each data point represents the mean +/- SEM. Statistical analyses for SRB growth curves were performed by completing a one-way ANOVA at the latest time point.
